# Rhythmic variations in prefrontal inter-neuronal correlations, their underlying mechanisms and their behavioral correlates

**DOI:** 10.1101/784850

**Authors:** S. Ben Hadj Hassen, C. Gaillard, E. Astrand, C. Wardak, S. Ben Hamed

## Abstract

Functional neuronal correlations between pairs of neurons are thought to play an important role in neuronal information processing and optimal neuronal computations during attention, perception, decision-making and learning. These noise correlations are often assumed to be stable in time. However, recent studies suggest that cognitive processes are rhythmic, this rhythmicity accounting for variations in overt behavioral performance. Whether this rhythmicity coincides with variations in shared noise variability is unknown. Here, we perform simultaneous recordings from the macaque frontal eye fields, while animals are engaged in a spatial memory task. We report that noise correlations in prefrontal cortex fluctuate rhythmically in the high alpha (10-16Hz) and beta (20-30Hz) frequency ranges. Importantly, these rhythmic modulations in shared neuronal variability account for dynamic changes in overt behavioral performance. They also coincide with increased spike-LFP phase coupling in these specific frequency ranges, the spatial profile of which vary between superficial and deep cortical layers. Finally, we demonstrate, using an artificial neuronal model, that rhythmic variations in noise correlation oscillations parsimoniously arise from long range (LFP) and local spike-LFP phase coupling mechanisms. Thus a significant portion of noise correlation fluctuations can be attributed to long-range global network rhythmicity.

## Introduction

Neuronal responses to the same stimulus fluctuate in time and across repetitions ^1,2^. This response variability is thought to be shared among functionally close neurons and is often referred to as noise correlations ^3^. These noise correlations reflect the amount of co-variability, in the trial-to-trial fluctuations in the response of pairs of neurons, to repeated presentations of identical stimuli, or under identical behavioral conditions, in the absence of any sensory stimulation. However, the exact origin of this shared neuronal variability remains unclear. It has been proposed that it arises from shared connectivity ^4^, global fluctuations in the excitability of cortical circuits ^5,6^, feedback signals ^7^, internal areal dynamics ^8–10^, as well as bottom-up peripheral sensory processing ^11^.

In fact, noise correlations have received a lot of attention and have been measured in a variety of brains areas, under numerous behavioral and stimulus conditions. Several studies suggest that noise correlations have a critical impact on cortical signal processing as well as onto behavioral performance ^3,11,11–14^, with learning or changes in behavioral state and attention ^2,15–23^.

The majority of these studies have measured how noise correlations are affected by spatial attention orientation, assuming stability in time. However, recent studies suggest that cognitive processes are based on rhythmic mechanisms that take place in the theta and alpha frequency ranges. This rhythmicity accounts for variations in overt behavioral performance ^24,25^. Whether this rhythmicity coincides with variations in shared noise variability is unknown. ^26^ describe variations in shared noise variability in the gamma band. Here, we demonstrate variations in prefrontal noise correlations in the alpha and beta frequency ranges. To achieve this, we recorded neuronal responses from macaque frontal eye fields (FEF), a cortical region at the source of spatial attention control signals ^27–30^ and in which noise correlations have been shown to vary as a function of spatial attention ^15^ and spatial memory ^31,32^. Monkeys were engaged in a spatial memory task. Overall, we demonstrate for the first time, rhythmic modulations of prefrontal noise correlations in two specific functional frequency ranges: the high alpha (10-16Hz) and the beta (20-30Hz) frequency ranges. Crucially, we show that these rhythmic modulations in noise correlations account both for overt behavioral performance and for layer specific modulations in spike-field phase coupling. Based on an artificial model, we demonstrate that rhythmic variations in noise correlation oscillations parsimoniously arise from long range (LFP) and local spike-LFP phase coupling mechanisms.

## Results

Neuronal recordings were performed in the prefrontal cortex, specifically in the frontal eye field (FEF, figure 1A), a structure known to play a key role in covert spatial attention ^28,33–35^. In each session, multi-unit activity (MUA) and local field potential (LFP) were recorded bilaterally, while monkeys performed a memory guided saccade task (figure 1B). Specifically, monkeys were required to hold the position of a spatial cue in memory for 700 to 1900ms and to perform a saccade towards the memorized spatial location on the extinction of the fixation point that served as a go signal. In the following, noise correlations between the different prefrontal signals of the same hemisphere were computed during the time interval running from 300ms to 1500ms following cue offset, on neuronal activities averaged over 200ms sliding windows (step of 10ms). As shown by previous studies, noise correlations decrease as a function of cortical distance (Figure S1A, 1-way ANOVA, p<0.001, Wilcoxon rank sum test, p<0.001 for 750 µm, p<0.001 for 1000 µm, ^23,36,37^ and are significantly lower among neuronal pairs with different spatial selectivity than neuronal pairs with the same spatial selectivity (Figure S1B, 1-way ANOVA, p<0.001)^38^.

**Figure 1:**
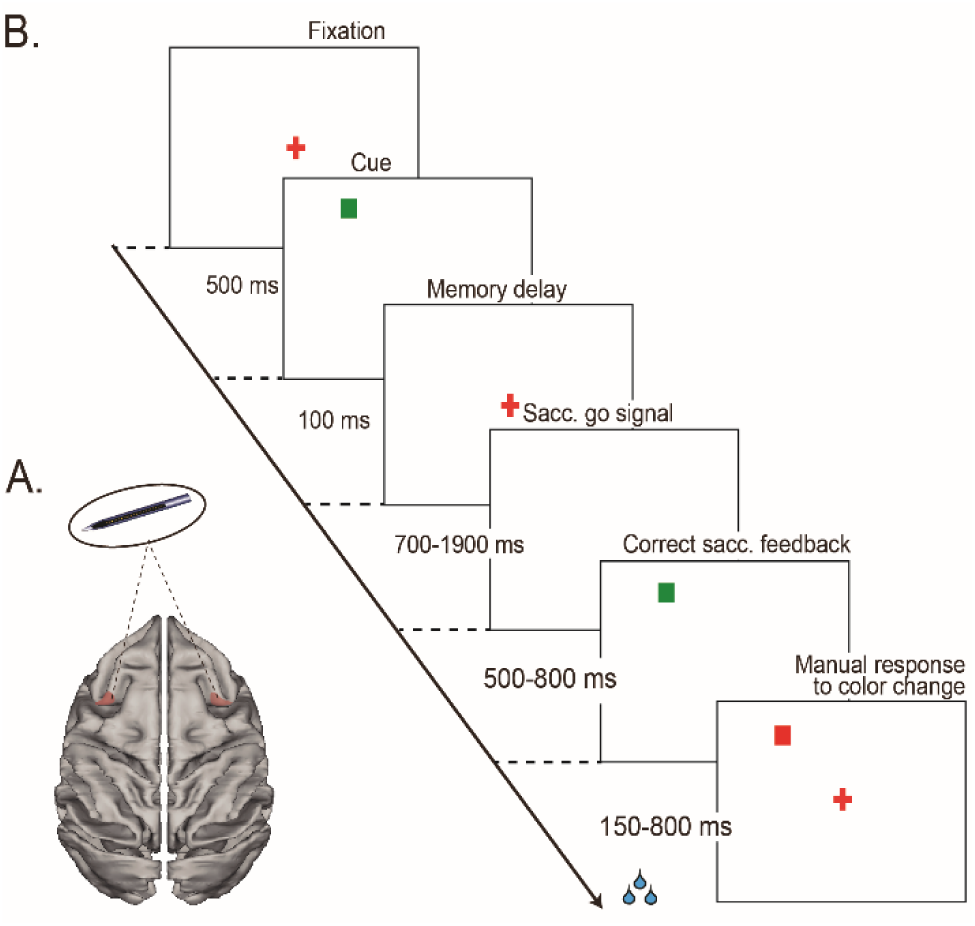
***(A) Recordings sites.*** On each session, 24-contact recording probes were placed in the left and right FEFs. ***(B) Memory-guided saccade task***. Monkeys had to fixate a red central cross. A visual cue was briefly flashed in one of four possible locations on the screen. Monkeys were required to hold fixation until the fixation cross disappeared and then produce a saccade to the spatial location indicated by the cue fixation point offset. On success, the cue re-appeared and the monkeys had to fixate it. They were then rewarded for producing a manual response 150ms to 800ms following the color change of this new fixation stimulus.

### Rhythmic fluctuations in noise correlations modulate behavioral response

Very few studies have addressed the question of how noise correlations vary as a function of time. ^26^ show that in primary visual cortex V1, noise correlations between neurons are modulated by gamma phase, synchronization in the 35-60Hz gamma-band producing maximal stimulus selectivity as well as minimal noise correlations. Whether this generalizes to other cortical regions and whether these variations in noise correlations are of behavioral relevance is currently unknown.

Here, we quantify variations in noise correlations during the cue to saccade go signal epoch, away from the initial sensory processing of the spatial cue. Specifically, in each session (n=26), noise correlations were computed between each pair of task-responsive channels (n=671, see Methods), during the spatial memory delay, running from 300ms to 1500ms following cue offset. During this epoch monkeys were required to memorize the cue location and get prepared to produce a spatially oriented saccade in response to an unexpected saccade go signal (fixation cross offset). In these computations, we included only trials with cue to go signal duration longer than 1500ms. Figure 2A shows clear noise correlation fluctuations in time during a representative recording session. Across all session, noise correlations were characterized, during the spatial memory delay, by rhythmic fluctuations taking place in two distinct frequency ranges: a high alpha frequency range (10-16 Hz) and a beta frequency range (20-30Hz), as quantified by a wavelet analysis (figure 2B). Overall, this indicates that noise correlations are rhythmic, their oscillatory pattern being probably reset by cue presentation as has been shown for other types of neuronal oscillatory patterns; ^25,39–43^.

**Figure 2:**
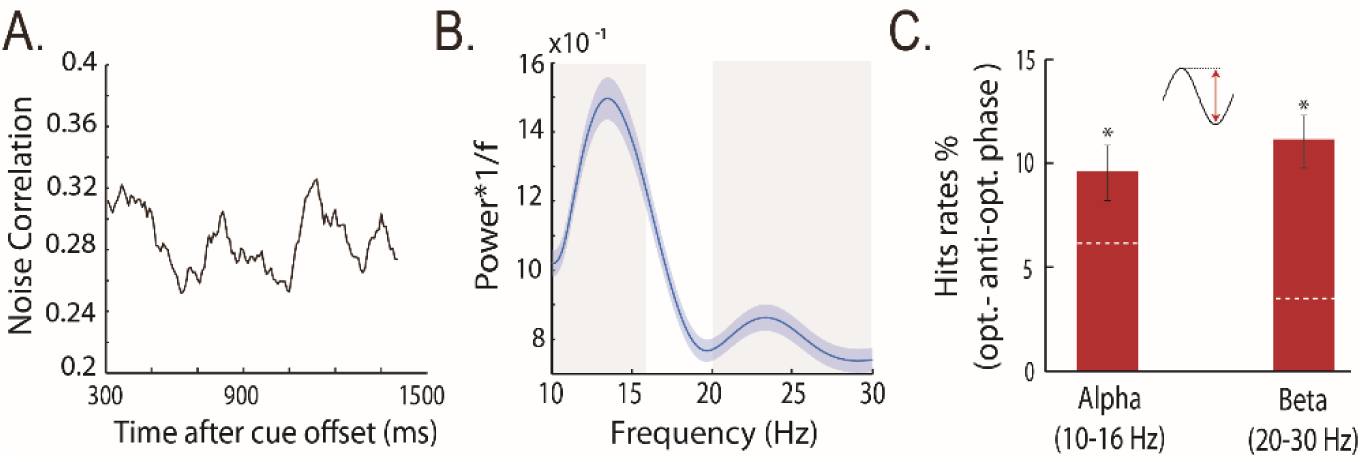
Rhythmic fluctuations in noise correlations modulate behavioral response. (A) Single memory guided saccade session example of noise correlation variations as a function of trial time. (B) 1/f weighted power frequency spectra of noise correlation in time (mean +/- s.e.), calculated from 300ms to 1500ms following cue offset. (C) Hit rate modulation by alpha and beta noise correlation at optimal phase as compared to anti-optimal phase, average +/- s.e., dashed white lines represent the 95% confidence interval under the assumption of absence of behavioral performance phase dependence.

An important question is whether these rhythmic variations in noise correlations contribute to task-related information processing and account for variations in overt behavioral performance. Indeed, low noise correlation neuronal population states are found to correlate with high neuronal population informational content ^3^ as well as with high behavioral performance states, i.e. correct trials as compared to incorrect trials ^44^. Here, we show that overt behavioral performance; defined as the proportion of correct trials as compared to misses trials, vary as a function of alpha and beta noise correlation oscillations. Specifically, on a session by session basis, we identify an optimal alpha (10-16Hz) phase for which the behavioral performance is maximal (+9.5%) compared to the corresponding anti-phase for which the behavioral performance is lowest (figure 2C). Similarly, an optimal beta (20-30Hz) phase is found to modulate behavioral performance in the same range of amplitudes (+11%). Overall, we thus demonstrate that the phases of alpha and beta oscillations in prefrontal noise correlations are predictive of overt behavioral performance.

### Oscillations in noise correlations coincide with enhanced spike-LFP phase coupling (SFC) in specific frequency ranges

High alpha and beta oscillations in the local field potentials (LFP) are ubiquitous and are considered to reflect long-range processes. Beta oscillations have been associated with cognitive control and cognitive flexibility. On the other hand, alpha oscillations are associated with attention, anticipation ^45,46^, perception ^47–49^, and working memory ^50^. A parsimonious hypothesis is thus that oscillations in noise correlations also arise from long-range process, via specific SFC mechanisms. Confirming this hypothesis, figure S2 represents SFC (as assessed from a PPC analysis, see Materials and Methods), computed during a 1200ms time interval over the spatial memory delay, running from 300ms to 1500ms following cue offset. Spike-LFP phase coupling peaks at the same frequency ranges identified in the noise correlation spectra, namely the high alpha range (10-16Hz) and the beta range (20-30Hz). When considering SFC independently of cortical layer organization, no difference in SFC is found between preferred and non-preferred positions in either frequency ranges. Interestingly, these selective SFC mechanisms are independent from overall LFP power content. Indeed, LFP power on the same dataset shows a deviation from the 1/f expect drop in a broad frequency range running from 15-30 Hz (figure S3A). This increase in LFP power is higher after cue presentation as compared to before (figure S3B). In contrast with SFC, this increase in LFP power is also more pronounced when the monkey is being cued towards the preferred than towards the non-preferred spatial location of the recorded signals (figure S3B). Overall, this thus suggests that oscillations in noise correlations arise from specific phase coupling mechanisms between long-range incoming LFP signals and local spiking mechanisms, independently from phase-amplitude coupling mechanisms.

### Spike-LFP phase coupling (SFC) differs between superficial and deep FEF layers

FEF neurons are characterized by a strong visual, saccadic, spatial memory and spatial attention selectivity ^27,28,51^. Previous studies have shown that pure visual neurons are predominantly located in the supragranular layers of the FEF while visuo-motor neurons are predominantly located in its infragranular layers ^51–56.57^ further show that supragranular FEF neurons predominantly project to striate visual cortex while infragranular FEF neurons predominantly project to the superior colliculus ^58–60^. The question we address here is whether the specific phase coupling mechanisms identified in the previous section are common to both supra- and infragranular FEF layers. Buffalo et al. have shown that, in extra-striate area V4, the ratio between the alpha and gamma spike field coherence discriminate between LFP signals in deep (low alpha / gamma spike field coherence ratio) and superficial cortical layers (high alpha / gamma spike field coherence ratio, ^61^. In our own data, as recordings were performed tangentially to FEF cortical surface, we have no direct assignation of the recorded MUAs to either superficial or deep cortical layers. However, the LFP alpha / gamma spike field coherence ratio provides a very reliable segregation of visual and visuo-motor MUAs at the same recording sites (figure 3A). We thus consider that, as has been described for area V4, this measure allows for a reliable delineation of superficial and deep layers in area FEF.

**Figure 3:**
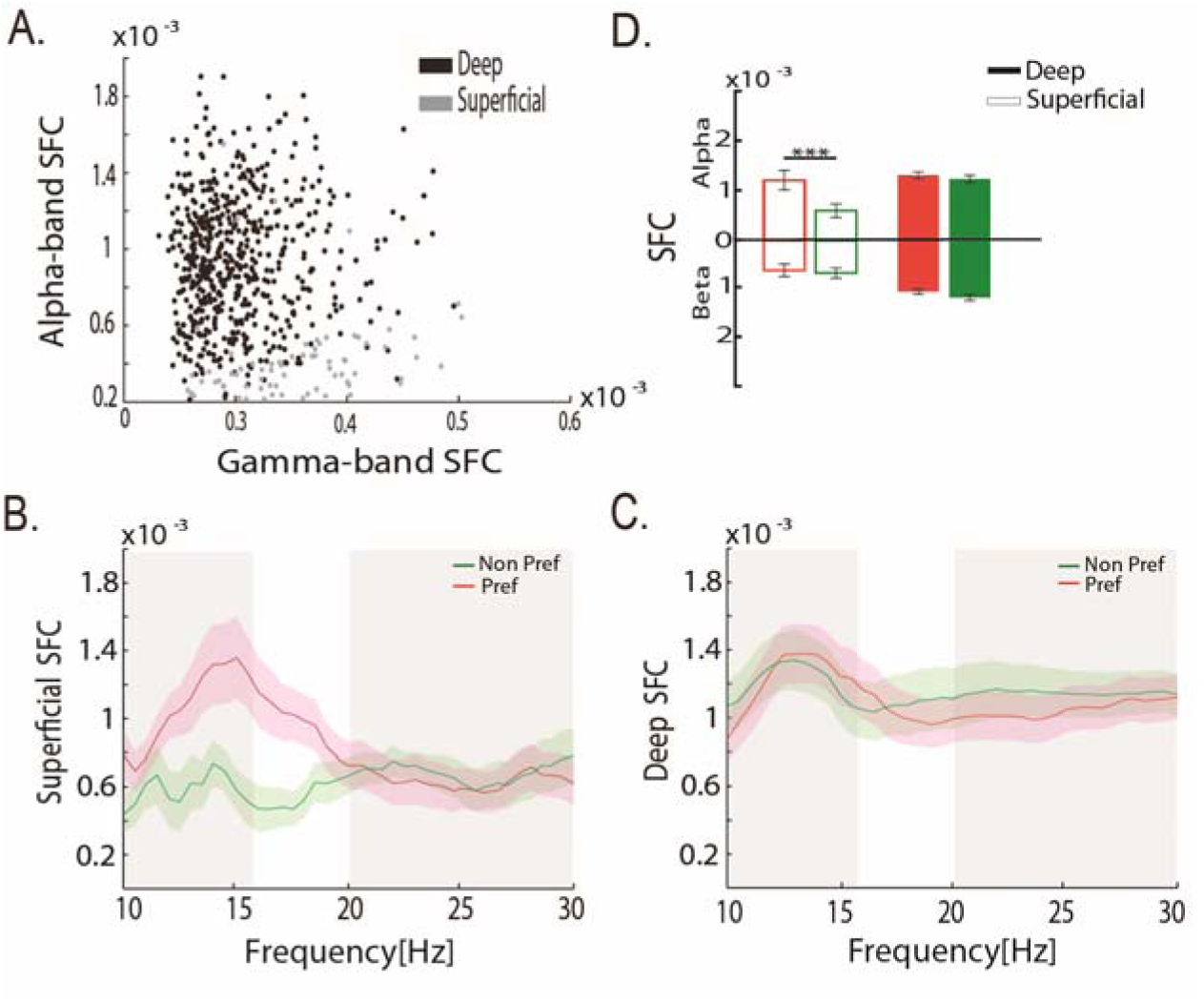
Spike-LFP phase coupling (SFC) and noise correlations differ between superficial and deep layers. (A) Distribution of SFC in gamma- and - alpha-bands for superficial and deep layers in area FEF. (B) Average SFC (mean +/- s.e. across sessions, calculated during 300ms to 1500ms following cue offset within superficial cortical layer signals. (C) Same as (B) but within deep cortical layer. (D) Average SFC (+/- s.e.) in alpha (10-16Hz, top histogram) and beta (20-30Hz, bottom histogram) for superficial (empty bars) and deep (filled bars) cortical layer signals (t-test, ***: p<0.001).

Figure 3B-C represents the SFC applied to the same data as presented in figure S3, but separated on the bases of the attribution of the MUA to either superficial or deep cortical FEF layers. While SFC modulations are observed in the same frequencies of interest as in figure S3, i.e. in the high alpha range (10-16Hz) and the beta range (20-30Hz), clear layer differences can be observed (figure 3B-C). Specifically, within superficial layers (figure 3B), SFC is selectively enhanced in the high alpha (10-16Hz) frequency range when spatial memory is oriented towards the preferred spatial location of the recorded signals as compared to away (figure 3D, p<0.001). This coincides with an enhanced high alpha power in the preferred compared to the non-preferred condition in the superficial layers (figure S4). No difference in SFC is observed in the beta frequency range, for preferred vs. non-preferred locations. In deep layers (figure 3C), SFC is enhanced in both the higher alpha and beta frequency ranges, irrespectively of whether spatial memory is oriented towards or away from the preferred spatial locations (figure 3D). This coincides, in the deep layers, with high alpha power in both the preferred and non-preferred conditions (figure S4). This result suggests distinct selective control mechanisms of correlated noise, spatially selective in superficial FEF layers and non-spatially selective in deep FEF layers.

### Modeling rhythmic variations in noise correlations

In this last section, using an artificial neuronal population reproducing observed spike and LFP parameters, we provide a causal and parsimonious model linking spike-field coherence and noise correlations mechanisms. The input data to the model are superficial and deep LFP signals generated to match the experimental observation of high alpha / gamma SFC ratio in superficial cortical layers and low alpha / gamma SFC ratio in deep cortical layers (figure S5) as well as FEF LFP power content as a function of preferred and non-preferred spatial memory (as per figures S3 and S4). Spike data are generated such that SFC is high in the high alpha and beta frequency ranges. In the model, differences in the input LFP power between the superficial and deep layers in the preferred and non-preferred spatial memory conditions combined with selective SFC in the high alpha and beta frequencies are sufficient to reproduce the empirical SFC differences (figure 4A and 4B to be compared to figure 3B and 3C). The resultant spiking population is characterized by variable noise correlations in time (figure 4C) with a marked rhythmic pattern in the high alpha and beta frequency ranges (figure 4D). Importantly, these rhythmic properties of neuronal noise correlations were resilient to changes in LFP power outside the SFC frequency ranges. Overall, this thus points towards a local origin for the reported noise correlation oscillations, implemented via selective SFC spiking mechanisms that differ in their spatial selectivity between superficial and deep layers. The output of this local mechanism is however further modulated by long-range influences captured by the low frequency alpha and beta LFP frequency content.

**Figure 4:**
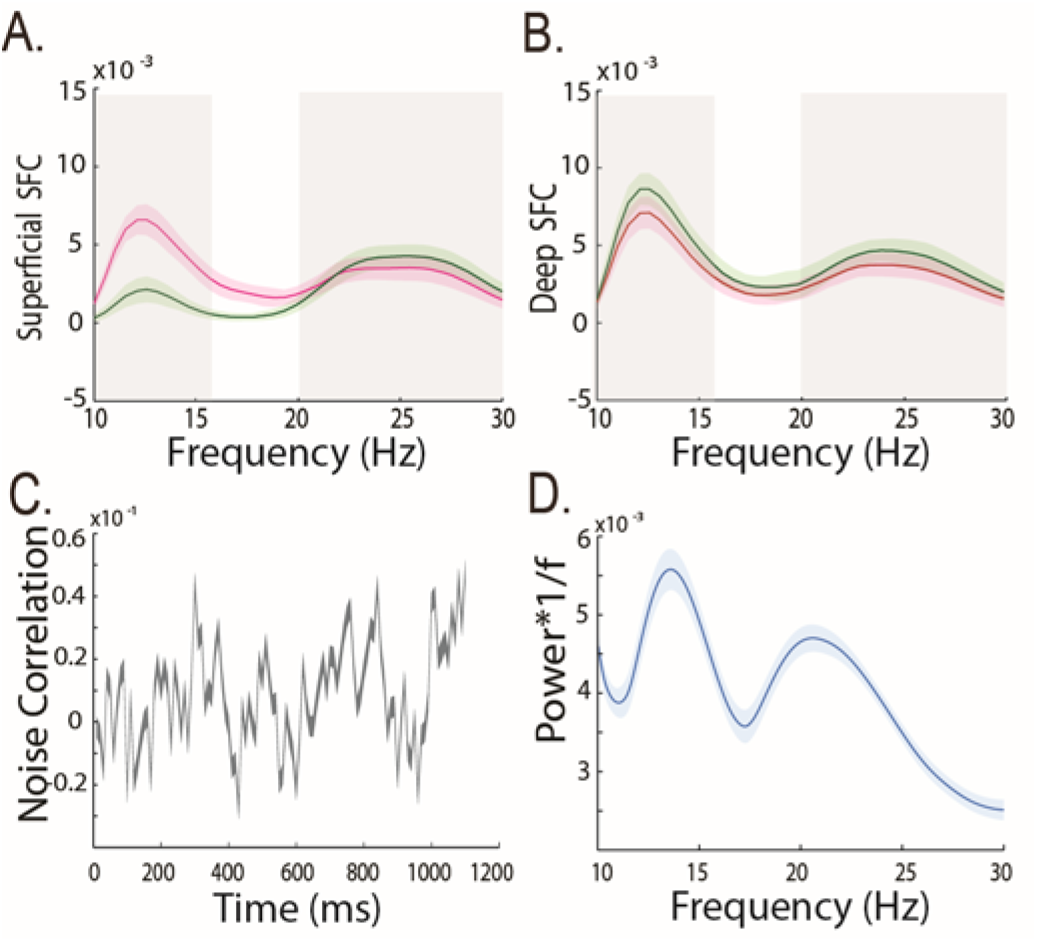
Modelling of rhythmic variations in noise correlations. (A) Model average SFC (mean +/- s.e.) within superficial cortical layer signals. (B) Same as in (A) but within deep cortical layer signals. (C) Model example of noise correlation variations as a function of trial time. (D) 1/f weighted power frequency spectrum of noise correlation in time (mean +/- s.e.) presented in (C).

Overall, we thus propose a model that shows that the observed rhythmic variations in noise correlations can be parsimoniously explained by selective SFC mechanisms in the higher alpha and beta frequency ranges, the strength of which are modulated by LFP power in these specific frequency bands irrespective of other frequencies.

## Discussion

### Oscillations in noise correlations at multiple time scales

Here, we describe, in the prefrontal cortex, two distinct regimens of rhythmic fluctuations in noise correlations, namely in the high alpha (10-16 Hz) frequency range as well as in the beta (20-30Hz) frequency range. These dominate over faster fluctuations in the gamma band (data not shown), as described in the primary visual cortex by ^26^, and which have been shown to coincide with variations in stimulus selectivity and enhanced gamma-band synchronization. Interestingly, identical rhythmic fluctuations in noise correlations can also be identified in the parietal cortex (LIP, data not shown). FEF and LIP belong to the same functional network ^28,62^ and are densely interconnected ^63,64^. It is thus not surprising that both cortical areas share the same noise correlation rhythmic properties, and supports a long-range origin for these rhythmic patterns (see below). Whether this noise correlations rhythms are ubiquitous and extend to, for example, the primary visual cortex, or whether they are specific to the parieto-frontal cortex and in tight link with the role of this functional network in attentional processes remains to be explored ^28,62,65^. Importantly, these rhythmic fluctuations in noise correlations are not specific to spatial memory processes, and can be observed in simple fixation or target detection tasks ^66^.

### Rhythmic noise correlations in neuronal population impact information capacity and behavior

The information capacity of a population code is thought to decrease as correlated noise among neurons increases ^3,13,14,67^, thought recent studies suggest that this detrimental aspect depends of noise correlation sources ^12,68^. Accordingly, fluctuations in noise correlations levels are expected to coincide with fluctuations in neuronal information. In V1, gamma-band fluctuations in primary visual cortex noise correlations are associated with variations in stimulus selectivity ^26^. Under the legitimate assumption of a direct relationship between prefrontal neuronal population information content and subjects’ behavior, a strong prediction of the dependency between overall neuronal population information capacity and noise correlations is a link between noise correlations and overt behavior. Here, we show that overt behavioral performances co-vary with both the alpha and beta noise correlation oscillations, accounting for up to 10% of the behavioral response variability. This indicates a functional role for these alpha and beta oscillations in noise correlations and supports the idea that noise correlation is a flexible physiological parameter that modulates overall neuronal population information capacity. Recent studies show that noise correlation contributes to optimally meet ongoing behavioral demands, during learning and attention ^69^. In a twin study, we show that noise correlations vary in strength both as a function of the ongoing task as well as a function of the time in the task, thus adjusting dynamically to ongoing behavioral demands ^66^.

### Cognitive rhythms and noise correlations

Alpha oscillations are consistently associated with attention, anticipation ^45,46^, perception ^47–49^, and working memory ^50^. Beta oscillations are on the other hand consistently associated with cognitive control and cognitive flexibility ^62^. Gamma-oscillations reflect local neuronal processes propagating in the feedforward direction ^70^ and spatial attention orientation coincides with increased gamma oscillations ^33,71^ as well as increased SFC ^72,73^. In contrast, alpha ^70^ and beta ^62^ oscillations reflect long-range processes propagating in the feedback direction, and spatial attention orientation coincides with decreased alpha and beta SFC. Our recordings are performed in the FEF, a cortical region which has been demonstrated to be at the source of spatial attention control signals ^27–30^. In this cortical region, rhythmic processes in relation with spatial attention deployment have recently been described both in the theta ^25,74^ and in the lower alpha ^83^ frequencies, supporting the hypothesis that attention is an intrinsically rhythmic cognitive process ^25^. The oscillations in noise correlation described here take place in frequency ranges that are independent from those described in spatial attention and memory studies. This suggests that they are of a different neuronal origin and correlate with neuronal mechanisms that are distinct from those at play during selective spatial cognitive processes.

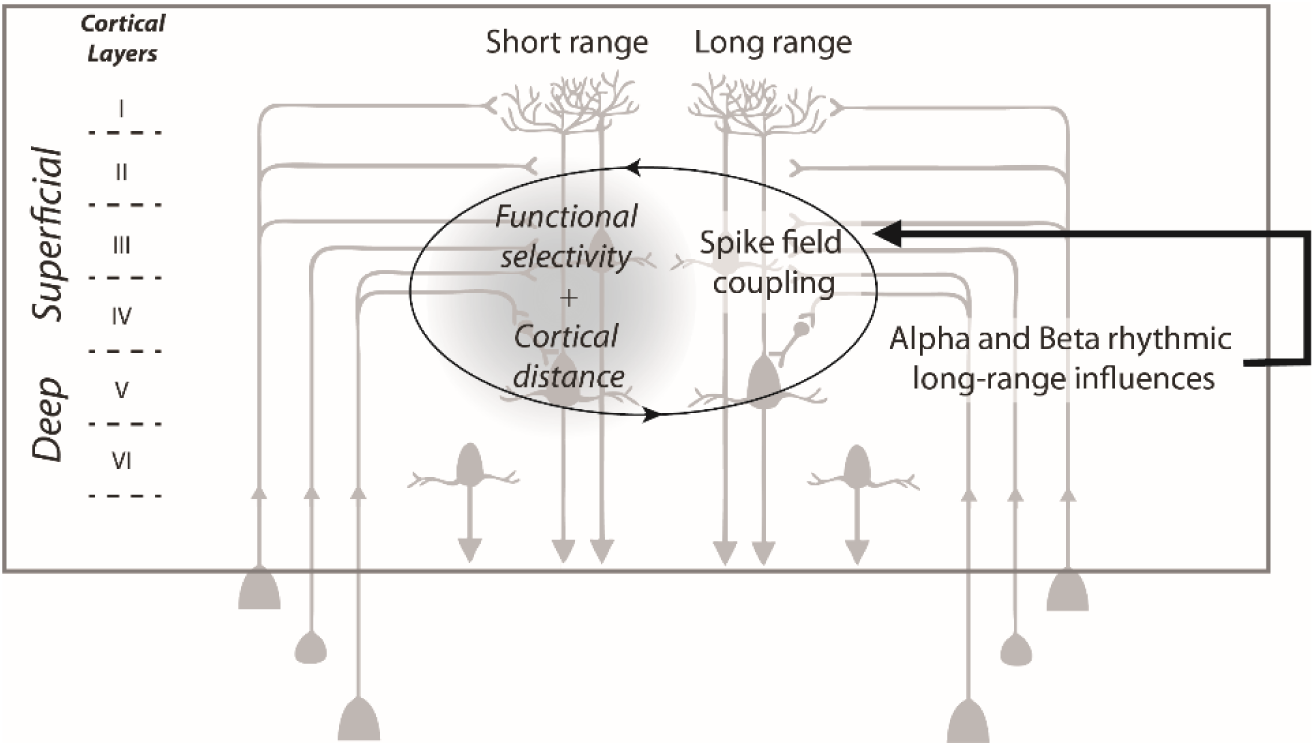
Sources of noise correlations. Noise correlations are shaped by short-range inter-neuronal connectivity that depend both on cortical distance and on functional selectivity (left) and by alpha and beta rhythmic long-range influences that differentially impact spike LFP phase coupling in the superficial and deep cortical layer.

### Mechanism accounting for rhythmic noise correlations

The rhythmic fluctuations in noise correlations we describe here co-exist with rhythmic fluctuations of SFC in the same frequency ranges. This supports a functional link between these two processes. The origin of this shared noise variability is still a subject of debate; it can arise from the afferent pathways ^2,4^, from top-down signals ^75,76^, or from coherent synchronization mechanisms in functional sub-networks. Our model confirms that rhythmic variations in noise correlations are parsimoniously accounted for by joint long-range influences reflected in the LFP alpha and beta ranges and selective local SFC mechanisms in these very same frequencies. This causal link being irrespective of changes in LFP power away from these frequency ranges. In other words, long-range high alpha and beta modulate the degree of synchronization between local neuronal populations (figure 5, right). Confirming this observation, we show that at the same time as the strength of noise correlation high alpha and beta oscillations vary as a function of the ongoing task, so does SFC ^66^. Supporting these long-range influences on noise correlations, we show that, in the absence of spatial memory signals, SFC modulation in the alpha range strongly decreases in the more superficial cortical layers as compared to the deeper layers. We propose that SFC coupling selectivity to specific frequency ranges is due to the biophysical membrane properties of specific prefrontal cell types ^77^. This will need to be further investigated. In addition to these long-range modulations, local recurrent connectivity also affects noise correlations ^12^. This results in the classical observation that shared neuronal variability decreases as a function of cortical distance as well as a function of functional dissimilarity between neuronal pairs (figure 5, left). The extent to which these local recurrent mechanisms differ between superficial and deeper layers remains to be explored.

Overall, this thus leads to the strong prediction that local flexibility in noise correlations as a function of behavioral demand arises from changes in long-range incoming signals in specific frequency bands, namely the high alpha and beta frequencies, and acts independently from other previously described neuronal processes such as spatial attention and spatial memory processes. The exact source of these signals, their relation to behavioral optimization and flexibility and how they interact and are integrated with other sources of variations in noise correlations remain to be explored.

## Acknowledgments

S.B.H.H. was supported by ANR grant ANR-14-CE13-0005-1. C.G. was supported by the French Ministère de l’Enseignement Supérieur et de la Recherche. S.B.H. was supported by ANR grant ANR-11-BSV4-0011, ANR grant ANR-14-CE13-0005-1, and the LABEX CORTEX (ANR-11-LABX-0042) of Université de Lyon, within the program Investissements d’Avenir (ANR-11-IDEX-0007) operated by the French National Research Agency (ANR). E.A. was supported by the CNRS-DGA and Fondation pour la Recherche Médicale. We thank research engineer Serge Pinède for technical support and Jean-Luc Charieau and Fabrice Hérant for animal care. All procedures were approved by the local animal care committee (C2EA42-13-02-0401-01) in compliance with the European Community Council, Directive 2010/63/UE on Animal Care.

## Authors contributions

Conceptualization, S.B.H. S.B.H.H. and C.G.; Methodology, S.B.H., S.B.H.H., C.G., E.A., C.W.; Investigation, S.B.H., S.B.H.H., C.G., E.A. and C.W.; Writing – Original Draft, S.B.H. S.B.H.H. and C.G.; Writing – Review & Editing, S.B.H. S.B.H.H. and C.G.; Funding Acquisition, S.B.H.; Supervision, S.B.H.

## Supplementary material and methods

### Material and methods

#### Ethical statement

All procedures were in compliance with the guidelines of European Community on animal care (Directive 2010/63/UE of the European Parliament and the Council of 22 September 2010 on the protection of animals used for scientific purposes) and authorized by the French Committee on the Ethics of Experiments in Animals (C2EA) CELYNE registered at the national level as C2EA number 42 (protocole C2EA42-13-02-0401-01).

#### Surgical procedure

As in^44^, two male rhesus monkeys (Macaca mulatta) weighing between 6-8 kg underwent a unique surgery during which they were implanted with two MRI compatible PEEK recording chambers placed over the left and the right FEF hemispheres respectively (figure 1A), as well as a head fixation post. Gas anesthesia was carried out using Vet-Flurane, 0.5 – 2% (Isofluranum 100%) following an induction with Zolétil 100 (Tiletamine at 50mg/ml, 15mg/kg and Zolazepam, at 50mg/ml, 15mg/kg). Post-surgery pain was controlled with a morphine pain-killer (Buprecare, buprenorphine at 0.3mg/ml, 0.01mg/kg), 3 injections at 6 hours interval (first injection at the beginning of the surgery) and a full antibiotic coverage was provided with Baytril 5% (a long action large spectrum antibiotic, Enrofloxacin 0.5mg/ml) at 2.5mg/kg, one injection during the surgery and thereafter one each day during 10 days. A 0.6mm isomorphic anatomical MRI scan was acquired post surgically on a 1.5T Siemens Sonata MRI scanner, while a high-contrast oil filled grid (mesh of holes at a resolution of 1mmx1mm) was placed in each recording chamber, in the same orientation as the final recording grid. This allowed a precise localization of the arcuate sulcus and surrounding gray matter underneath each of the recording chambers. The FEF was defined as the anterior bank of the arcuate sulcus and we specifically targeted those sites in which a significant visual and/or oculomotor activity was observed during a memory guided saccade task at 10 to 15° of eccentricity from the fixation point (figure 1A). In order to maximize task-related neuronal information at each of the 24-contacts of the recording probes, we only recorded from sites with task-related activity observed continuously over at least 3 mm of depth.

#### Behavioral task

During a given experimental session, the monkeys were placed in front of a computer screen (1920×1200 pixels and a refresh rate of 60 Hz) with their head fixed. Their water intake was controlled so that their initial daily intake was covered by their performance in the task, on a trial by trial basis. This quantity was complemented as follows. On good performance sessions, monkeys received fruit and water complements. On bad performance sessions, water complements were provided at a distance from the end of the session. During a ***Memory-guided saccade Task*** (figure 1B): A red fixation cross (0.7×0.7°) appeared in the center of the screen and the monkeys were required to hold fixation for 500 msec, within a fixation window of 1.5×1.5°. A square green cue (0.28×0.28°) was then flashed for 100ms at one of four possible locations ((10°, 10°), (−10°, 10°), (−10°,-10°) and (10°,-10°)). The monkeys had to continue maintain fixation on the central fixation point for another 700–1900 ms until the fixation point disappeared. The monkeys were then required to make a saccade towards the memorized location of the cue within 500-800ms from fixation point disappearance, and a spatial tolerance of 4°x4°. On success, a target, identical to the cue was presented at the cued location and the monkeys were required to fixate it and detect a change in its color by a bar release within 150-800 ms from color change. Success in all of these successive requirements conditioned reward delivery.

#### Neural recordings

On each session, bilateral simultaneous recordings in the two FEFs were carried out using two 24-contact Plexon U-probes. The contacts had an interspacing distance of 250 µm. Neural data was acquired with the Plexon Omniplex® neuronal data acquisition system. The data was amplified 400 times and digitized at 40,000 Hz. The MUA neuronal data was high-pass filtered at 300 Hz. The LFP neuronal data was filtered between 0.5 and 300 Hz. In the present paper, all analyses are performed on the multi-unit activity recorded on each of the 48 recording contacts. A threshold defining the multi-unit activity was applied independently for each recording contact and before the actual task-related recordings started. All further analyses of the data were performed in Matlab™ and using FieldTrip ^78^ and the Wavelet Coherence Matlab Toolbox ^79^, both open source Matlab™ toolboxes.

#### Data Analysis

##### Data preprocessing

Overall, MUA recordings were collected from 48 recording channels on 26 independent recording sessions (13 for M1 and 13 for M2). We excluded from subsequent analyses all channels with less than 5 spikes per seconds. For each session, we identified the task-related channels based on a statistical change (one-way ANOVA, p<0.05) in the MUA neuronal activity in the memory-guided saccade task, in response to either cue presentation ([0 400] ms after cue onset) against a pre-cue baseline ([-100 0] ms relative to cue onset), or to saccade execution go signal and to saccade execution (i.e. fixation point off, [0 400] ms after go signal) against a pre-go signal baseline ([-100 0] ms relative to go signal), irrespective of the spatial configuration of the trial. In total, 671 channels were retained for further analyses out of 1248 channels.

##### Distance between recording sites

For each electrode, pairs of MUA recordings were classified along four possible distance categories: D1, spacing of 250 µm; D2, spacing of 500 µm; D3, spacing of 750 µm and D4, spacing of 1mm. These distances are an indirect proxy to actual cortical distance, as the recordings were performed tangentially to cortical surface, i.e. more or less parallel to sulcal surface.

##### MUA spatial selectivity

FEF neurons are characterized by a strong visual, saccadic, spatial memory and spatial attention selectivity^27,28,51^. We used a one-way ANOVA (p<0.05) to identify the spatially selective channels in response to cue presentation ([0 400] ms following cue onset) and to the saccade execution go signal ([0 400] ms following go signal). Post-hoc t-tests served to further order, for each channels, the neuron’s response in each visual quadrant from preferred (p1), to least preferred (p4). By convention, positive modulations were considered as preferred and negative modulations as least preferred. For example, in a given session, the MUA signal recorded on channel 1 of a probe placed in the left FEF, could have as best preferred position p1 the upper right quadrant, the next best preferred position p2 the lower right quadrant, the next preferred position p3 the upper left quadrant and the least preferred position p4 the lower left quadrant. The MUA signal recorded on channel 14 of this same probe, could have as best preferred position p1 the lower right quadrant, the next best preferred position p2 the upper right quadrant, the next preferred position p3 the lower left quadrant and the least preferred position p4 the upper left quadrant. Positions with no significant modulation in any task epoch were labeled as p0 (no selectivity for this position). Once this was done, for each electrode, pairs of MUA recordings were classified along two possible functional categories: pairs with the same spatial selectivity (SSS pairs, sharing the same p1) and pairs with different spatial selectivity (DSS pairs, such that the p1 of one MUA is a p0 for the other MUA). For the sake of clarity, we do not consider partial spatial selectivity pairs (such that the p1 of one MUA is a non-preferred, p2, p3 or p4 for the other MUA).

##### MUA layer attribution

As stated above, our recordings are not tangential to cortical surface. As a proxy to attribute a given recording channel to upper or lower cortical layers we proceeded as follows. For each electrode and each channel, we estimated, at the time of cue onset in the memory-guided saccade task (100ms-500ms from cue onset), the SFC in the alpha range (6 to 16 Hz) and the gamma range (40 to 60 Hz). Based on previous literature ^80^, we used the ratio between the alpha and gamma spike field-coherence as a proxy to assign the considered LFP signals to a deep cortical layer site (high alpha / gamma spike-field coherence ratio) or to a superficial cortical layer site (low alpha / gamma SFC ratio). We also categorized MUA signals into visual, visuo-motor and motor categories, as in ^81^. Briefly, average firing rates were computed in 3 epochs: [-100 0] ms before cue onset (baseline), [0 200] ms after cue onset (visual), and [0 200] ms before saccade onset (movement). Neurons with activity statistically significantly different from the baseline (Wilcoxon rank-sum test, *P* < 0.05) after cue onset were categorized as visual. Neurons with activity statistically significantly different from the baseline (Wilcoxon rank-sum test, *P* < 0.05) before saccade onset were categorized as oculomotor. Neurons that were active in both epochs were categorized as visuo-movement neurons. The LFP categorization along the alpha to gamma SFC ratio strongly coincided with the classification of the MUA signals into purely visual sites (low alpha and gamma SFC ratio, input FEF layers) and visuo-motor sites (high alpha and gamma SFC ratio, output FEF layers, figure 3).

##### Oscillations in noise correlations

To measure oscillatory patterns in the noise correlation time-series data, we computed for each session (N=12) noise correlations over time (over successive 200ms intervals, sliding by 10ms, running from 300ms to 1500ms following cue offset). Specifically, for each channel i, and each trial k, the average neuronal response r_i_(k) for the 200 ms interval was calculated and z-score normalized into z_i_(k), where z_i_(k)=r_i_(k)-µ_i_/std_i_ and µ_i_ and std_i_ respectively correspond to the mean firing rate and standard deviation around this mean during the interval of interest of the channel of interest i. This z-score normalization allows to capture the changes in neuronal response variability independently of changes in mean firing rates. Noise correlations between pairs of MUA signals during the interval of interest were then defined as the Pearson correlation coefficient between the z-scored individual trial neuronal responses of each MUA signal over all trials. Only positive significant noise correlations are considered, unless stated otherwise. In any given recording session, noise correlations were calculated between MUA signals recorded from the same electrode, thus specifically targeting intra-cortical correlations. Once noise correlation is calculated over time a wavelet transform ^78^ was then applied on each session’s noise correlation time series. To control that these tmporal noise correlation oscillations cannot be attributed to changes in spiking activity, a wavelet analysis was also run onto MUA time series data (not shown).

##### Modulation of behavioral performance by alpha and beta noise correlation phase

To quantify the effect of noise correlation oscillations onto behavioral performance, we used a complex wavelet transform analysis (Fieldtrip, Oostenveld et al. 2011) to compute, for each session, in the noise correlations, the phase of the frequencies of interest (alpha / beta) following cue offset. For each session, we identified hit and miss trials falling at zero phase of the frequency of interest (+/- π /140) with respect to target presentation. Hit rates (HR) were computed for this zero phase bin. We then shifted this phase window by π /70 steps and recalculated the HR, repeating this procedure to generate phase-detection HR functions, across all phases, for each frequency of interest ^82^. For each session, the phase bin for which hit rate was maximal was considered as the optimal phase. The effect of a given frequency (alpha or beta) onto behavior corresponds to the difference between HR at this optimal phase and HR at the anti-optimal phase (optimal phase + π). To test for statistical significance, observed hit/miss phases were randomized across trials so as to shuffle the temporal relationship between phases and behavioral performance. This procedure was repeated 1000 times. 95% CI was then computed and compared to the observed behavioral data.

##### Spikes-LFP PPC

For each selected channel, spikes-LFP phase coupling spectra (SCF) were calculated between the spiking activity obtained in one channel and the LFP activity from the next adjacent channel in the time interval running from 300ms to 1500ms following cue offset. We used a single Hanning taper and applied convolution transform to the Hanning-tapered trials. We equalized the number of trials for all conditions so as to prevent any bias that could be introduced by unequal numbers of trials. We used a 4 cycles length per frequency. The memory guided saccade task is known to involve spatial processes during the cue to target interval that bias spike field coherence. Thus, spikes-LFP phase coupling was measured separately for trials in which the cued location matched the preferred spatial location of the channel and trials in which the cued location did not match the preferred spatial location of the channel. Statistics were computed across channels x sessions, using a non-parametric Friedman test.

##### Model

The objective of this model is to test whether rhythmic variations in noise correlations can parsimoniously be explained by joint long-range influences reflected in the LFP alpha and beta ranges and selective local spike-LFP phase coupling mechanisms in these same frequencies. This was investigated through synthetic neuronal population activities the main features of which were parsimoniously driven by FEF recorded data as follows. Spikes/LFPs signals were generated to create a 200 channels X 100 trials X 1000 ms, structure. LFP signals were constructed from a noise frequency component following a 1/f power/frequency law, and a signal component ranging from 10Hz to 60Hz. To mimic our empirical data (figure S3B), superficial LFPs to a preferred spatial memory location were enriched in alpha power. Spiking activities were composed of a noise component (spikes being extracted from a random binary process) and a component locked to LFP alpha (6-16Hz) and beta (20-30Hz) frequency phases, thus resulting in high spikes-LFP phase coupling in these specific frequencies. The strength of the SFC in each of these frequency ranges was manipulated to reproduce the laminar differences observed experimentally between superficial and deep cortical layer. Gamma (40-60Hz) SFC was also enhanced in the superficial FEF layers to match our empirical data as well as previous reports from other cortical regions ^80^. It is to be noted that, by definition, the strength of SFC at a specific frequency is exclusively modulated by the LFP power in the same frequency range. In other words, LFP frequencies that are not phase locked to the spiking activity do not contribute to the SFC measure. This can easily be modeled (data not shown). Last, the functional selectivity of the synthetic channels (preferred/ non preferred) was mimicked by a 15% increase in firing rate in the preferred condition, while the non-preferred condition remained unchanged, this for both superficial and deep channels. Frequency and phase analyses were performed using Wavelet Transform Analyzes based on the Wavelet Coherence Matlab Toolbox, SFC analyzes were performed using adapted Fieldtrip toolbox functions (http://fieldtriptoolbox.org). The outcome of this phase-phase coupling analysis is independent of instantaneous spiking rates.

## Supplementary figures

**Figure S1:**
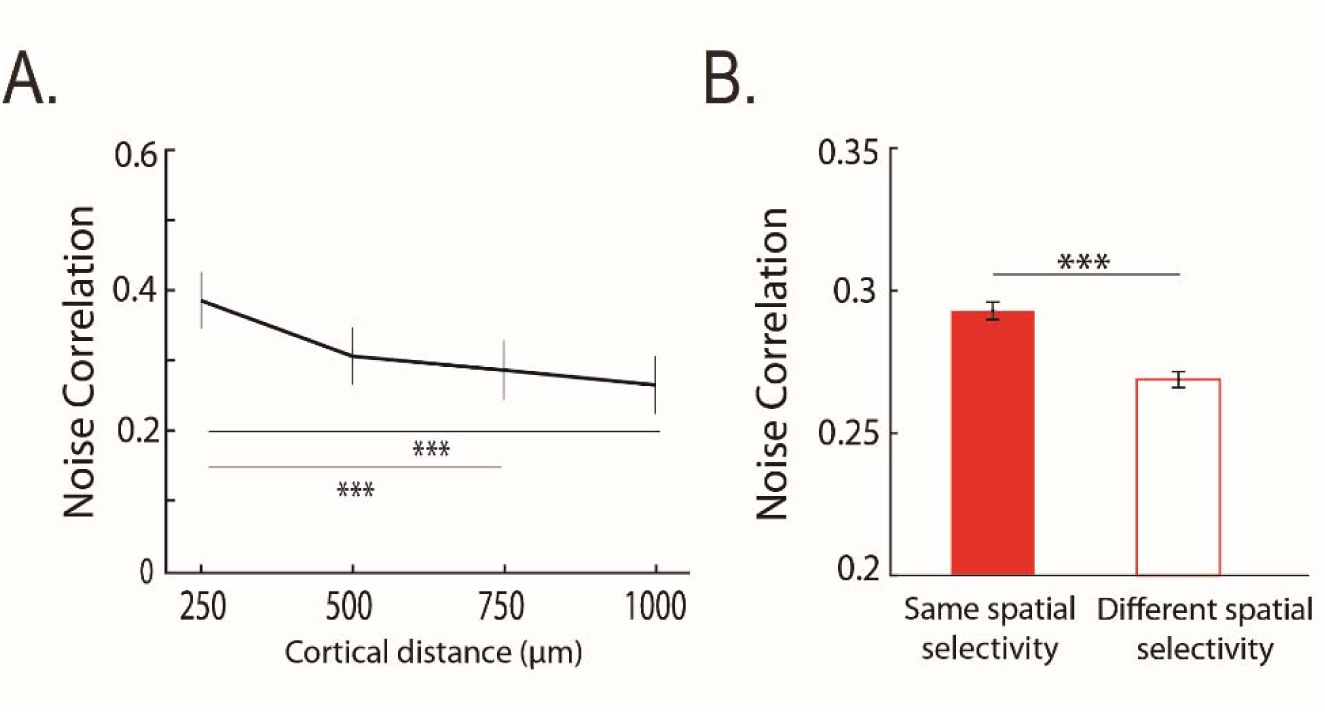
(A) Noise correlations as a function of cortical distance. Average noise correlations (mean +/- s.e.) across sessions, from 300 ms to 500ms after eye fixation onset, as a function of distance between pairs of channels: 250µm; 500µm; 750µm; 1000µm. (B) *Noise correlations as a function of spatial selectivity*. Average noise correlations (mean +/- s.e.) across sessions from 300ms to 500ms after eye fixation onset, as a function of whether noise correlations are calculated over signals sharing the same spatial selectivity (full bars) or not (empty bars). Stars indicate statistical significance following a two-way ANOVA and ranksum post-hoc tests; *p<0.05; **p<0.01; ***p<0.001.

**Figure S2:**
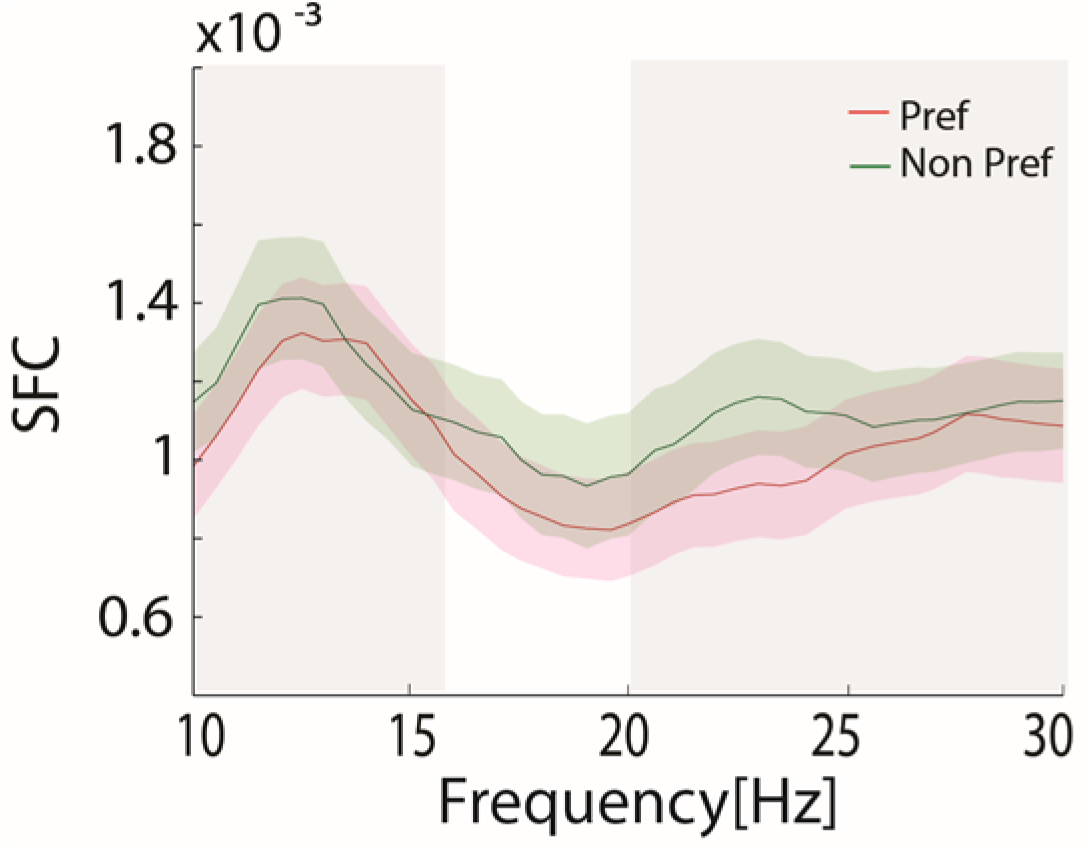
Average spikes-LFP phase coupling (mean +/- s.e.) across sessions, calculated during 300ms to 1500ms following cue offset as a function of preferred (red line) and non-preferred (green line) position.

**Figure S3:**
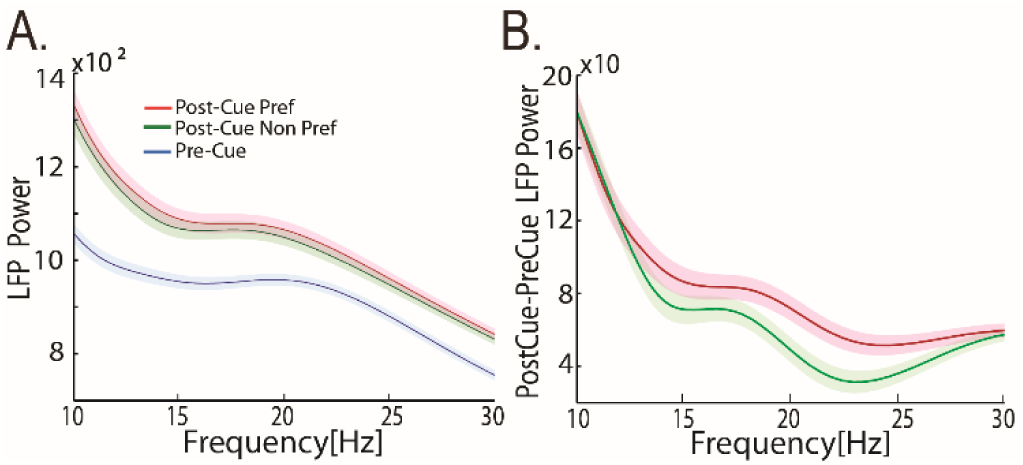
(A) Average of LFP power (mean +/- s.e.) across sessions, during pre-cue epoch (blue), post-cue epoch for the preferred position (red) and non-preferred position (green). (B) Difference of LFP power between post-cue preferred and non-preferred conditions for preferred (red) and non-preferred condition (green).

**Figure S4:**
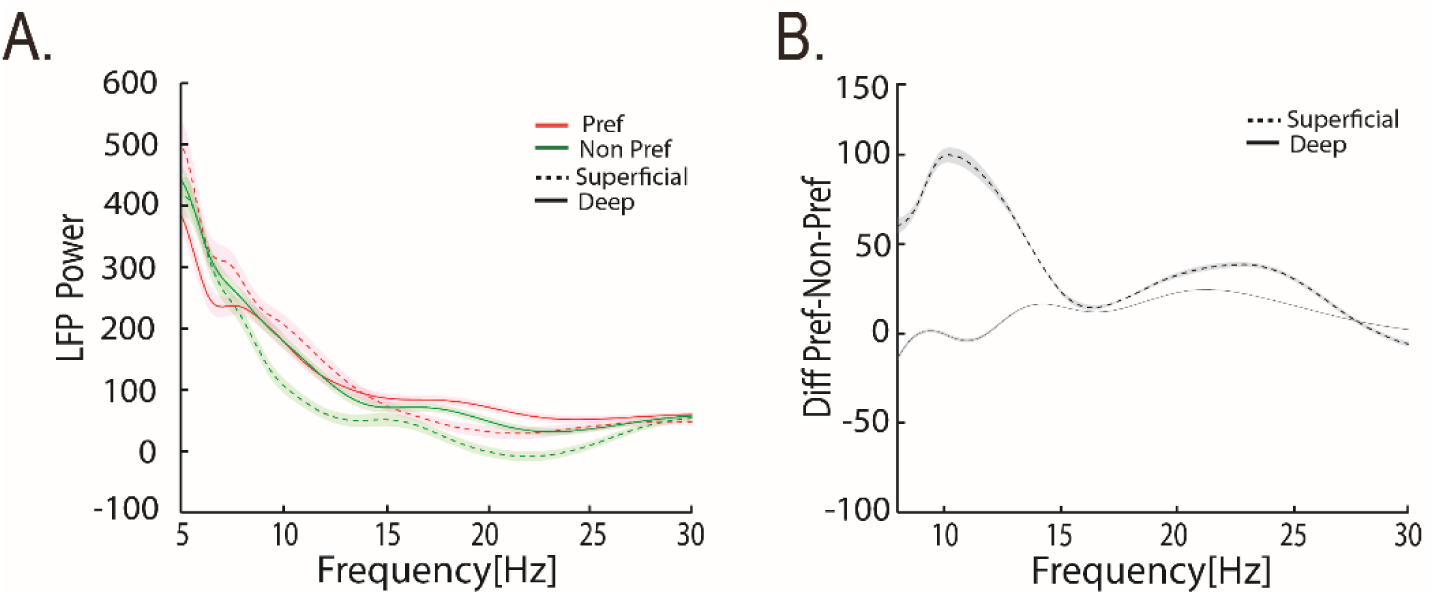
(A) Average of LFP power (mean +/- s.e.) across sessions, during post-cue epoch, for preferred position (red) and non-preferred position (green), in the superficial (dashed lines) and deep layers (continuous lines), relative to LFP power in pre-cue epoch. (B) Difference of LFP power between post-cue preferred and non-preferred conditions for superficial (dashed lines) and deep layers (continuous lines).

**Figure S5:**
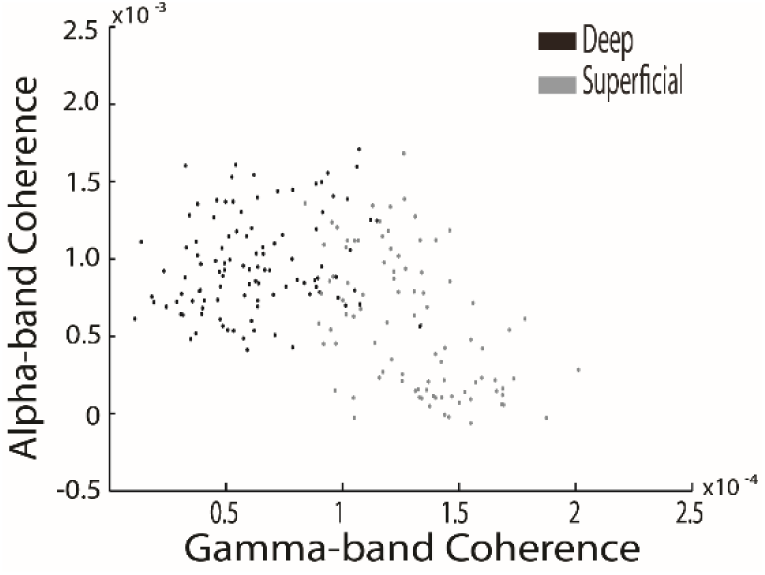
Model distribution of spikes-LFP phase coupling in gamma- and - alpha-bands for superficial and deep layers.

## References

1. Bach, M. & Kruger, J. Correlated neuronal variability in monkey visual cortex revealed by a multi-microelectrode. Exp. Brain Res. 61, (1986).

2. Zohary, E., Shadlen, M. N. & Newsome, W. T. Correlated neuronal discharge rate and its implications for psychophysical performance. Nature 370, 140–143 (1994).

3. Averbeck, B. B., Latham, P. E. & Pouget, A. Neural correlations, population coding and computation. Nat. Rev. Neurosci. 7, 358–366 (2006).

4. Shadlen, M. N. & Newsome, W. T. The variable discharge of cortical neurons: implications for connectivity, computation, and information coding. J. Neurosci. Off. J. Soc. Neurosci. 18, 3870–3896 (1998).

5. Ecker, A. S. et al. State Dependence of Noise Correlations in Macaque Primary Visual Cortex. Neuron 82, 235–248 (2014).

6. Goris, R. L. T., Movshon, J. A. & Simoncelli, E. P. Partitioning neuronal variability. Nat. Neurosci. 17, 858–865 (2014).

7. Wimmer, K. et al. Sensory integration dynamics in a hierarchical network explains choice probabilities in cortical area MT. Nat. Commun. 6, 6177 (2015).

8. Ben-Yishai, R., Bar-Or, R. L. & Sompolinsky, H. Theory of orientation tuning in visual cortex. Proc. Natl. Acad. Sci. U. S. A. 92, 3844–3848 (1995).

9. Litwin-Kumar, A. & Doiron, B. Slow dynamics and high variability in balanced cortical networks with clustered connections. Nat. Neurosci. 15, 1498–1505 (2012).

10. Ly, C., Middleton, J. W. & Doiron, B. Cellular and circuit mechanisms maintain low spike co-variability and enhance population coding in somatosensory cortex. Front. Comput. Neurosci. 6, 7 (2012).

11. Kanitscheider, I., Coen-Cagli, R. & Pouget, A. Origin of information-limiting noise correlations. Proc. Natl. Acad. Sci. U. S. A. 112, E6973–6982 (2015).

12. Moreno-Bote, R. et al. Information-limiting correlations. Nat. Neurosci. 17, 1410 (2014).

13. Sompolinsky, H., Yoon, H., Kang, K. & Shamir, M. Population coding in neuronal systems with correlated noise. Phys. Rev. E Stat. Nonlin. Soft Matter Phys. 64, 051904 (2001).

14. Abbott, L. F. & Dayan, P. The Effect of Correlated Variability on the Accuracy of a Population Code. (1999).

15. Cohen, M. R. & Maunsell, J. H. R. Attention improves performance primarily by reducing interneuronal correlations. Nat. Neurosci. 12, 1594–1600 (2009).

16. Gawne, T. J., Kjaer, T. W., Hertz, J. A. & Richmond, B. J. Adjacent visual cortical complex cells share about 20% of their stimulus-related information. Cereb. Cortex N. Y. N 1991 6, 482–489 (1996).

17. Gawne, T. J. & Richmond, B. J. How independent are the messages carried by adjacent inferior temporal cortical neurons? J. Neurosci. Off. J. Soc. Neurosci. 13, 2758–2771 (1993).

18. Gutnisky, D. A. & Dragoi, V. Adaptive coding of visual information in neural populations. Nature 452, 220–224 (2008).

19. Huang, X. & Lisberger, S. G. Noise Correlations in Cortical Area MT and Their Potential Impact on Trial-by-Trial Variation in the Direction and Speed of Smooth-Pursuit Eye Movements. J. Neurophysiol. 101, 3012–3030 (2009).

20. Mitchell, J. F., Sundberg, K. A. & Reynolds, J. H. Spatial attention decorrelates intrinsic activity fluctuations in macaque area V4. Neuron 63, 879–888 (2009).

21. Poort, J. & Roelfsema, P. R. Noise correlations have little influence on the coding of selective attention in area V1. Cereb. Cortex N. Y. N 1991 19, 543–553 (2009).

22. Reich, D. S. Independent and Redundant Information in Nearby Cortical Neurons. Science 294, 2566–2568 (2001).

23. Smith, M. A. & Kohn, A. Spatial and Temporal Scales of Neuronal Correlation in Primary Visual Cortex. J. Neurosci. 28, 12591–12603 (2008).

24. Fiebelkorn, I. C., Pinsk, M. A. & Kastner, S. The mediodorsal pulvinar coordinates the macaque fronto-parietal network during rhythmic spatial attention. Nat. Commun. 10, 215 (2019).

25. Fiebelkorn, I. C., Pinsk, M. A. & Kastner, S. A Dynamic Interplay within the Frontoparietal Network Underlies Rhythmic Spatial Attention. Neuron 99, 842-853.e8 (2018).

26. Womelsdorf, T. et al. Orientation selectivity and noise correlation in awake monkey area V1 are modulated by the gamma cycle. Proc. Natl. Acad. Sci. U. S. A. 109, 4302 (2012).

27. Astrand, E., Ibos, G., Duhamel, J.-R. & Ben Hamed, S. Differential dynamics of spatial attention, position, and color coding within the parietofrontal network. J. Neurosci. Off. J. Soc. Neurosci. 35, 3174–3189 (2015).

28. Ibos, G., Duhamel, J.-R. & Ben Hamed, S. A functional hierarchy within the parietofrontal network in stimulus selection and attention control. J. Neurosci. Off. J. Soc. Neurosci. 33, 8359–8369 (2013).

29. Moore, T., Armstrong, K. M. & Fallah, M. Visuomotor origins of covert spatial attention. Neuron 40, 671–683 (2003).

30. Wardak, C., Ibos, G., Duhamel, J.-R. & Olivier, E. Contribution of the monkey frontal eye field to covert visual attention. J. Neurosci. Off. J. Soc. Neurosci. 26, 4228–4235 (2006).

31. Meyers, E. M., Qi, X.-L. & Constantinidis, C. Incorporation of new information into prefrontal cortical activity after learning working memory tasks. Proc. Natl. Acad. Sci. U. S. A. 109, 4651–4656 (2012).

32. Constantinidis, C. & Klingberg, T. The neuroscience of working memory capacity and training. Nat. Rev. Neurosci. 17, 438–449 (2016).

33. Gregoriou, G. G., Gotts, S. J., Zhou, H. & Desimone, R. High-frequency, long-range coupling between prefrontal and visual cortex during attention. Science 324, 1207–1210 (2009).

34. Gregoriou, G. G., Gotts, S. J. & Desimone, R. Cell-type-specific synchronization of neural activity in FEF with V4 during attention. Neuron 73, 581–594 (2012).

35. Armstrong, K. M., Chang, M. H. & Moore, T. Selection and maintenance of spatial information by frontal eye field neurons. J. Neurosci. Off. J. Soc. Neurosci. 29, 15621–15629 (2009).

36. Constantinidis, C. & Goldman-Rakic, P. S. Correlated discharges among putative pyramidal neurons and interneurons in the primate prefrontal cortex. J. Neurophysiol. 88, 3487–3497 (2002).

37. Lee, D., Port, N. L., Kruse, W., Georgopoulos, A. P. & Neurology. Variability and correlated noise in the discharge of neurons in motor and parietal areas of the primate cortex. J Neurosci 18, 1161–1170. (1998).

38. Bair, W., Zohary, E. & Newsome, W. T. Correlated firing in macaque visual area MT: time scales and relationship to behavior. J. Neurosci. Off. J. Soc. Neurosci. 21, 1676–1697 (2001).

39. Dugué, L., Roberts, M. & Carrasco, M. Attention Reorients Periodically. Curr. Biol. CB 26, 1595–1601 (2016).

40. Fiebelkorn, I. C., Saalmann, Y. B. & Kastner, S. Rhythmic sampling within and between objects despite sustained attention at a cued location. Curr. Biol. CB 23, 2553–2558 (2013).

41. Landau, A. N. & Fries, P. Attention samples stimuli rhythmically. Curr. Biol. CB 22, 1000–1004 (2012).

42. VanRullen, R. Visual Attention: A Rhythmic Process? Curr. Biol. 23, R1110–R1112 (2013).

43. Corentin, G. et al. Prefrontal attentional saccades explore space at an alpha rhythm. bioRxiv 637975 (2019).

44. Astrand, E., Wardak, C., Baraduc, P. & Ben Hamed, S. Direct Two-Dimensional Access to the Spatial Location of Covert Attention in Macaque Prefrontal Cortex. Curr. Biol. CB 26, 1699–1704 (2016).

45. Thut, G., Nietzel, A., Brandt, S. A. & Pascual-Leone, A. Alpha-band electroencephalographic activity over occipital cortex indexes visuospatial attention bias and predicts visual target detection. J. Neurosci. Off. J. Soc. Neurosci. 26, 9494–9502 (2006).

46. Rihs, T. A., Michel, C. M. & Thut, G. A bias for posterior alpha-band power suppression versus enhancement during shifting versus maintenance of spatial attention. NeuroImage 44, 190–199 (2009).

47. Varela, F. J., Toro, A., John, E. R. & Schwartz, E. L. Perceptual framing and cortical alpha rhythm. Neuropsychologia 19, 675–686 (1981).

48. Mathewson, K. E., Gratton, G., Fabiani, M., Beck, D. M. & Ro, T. To See or Not to See: Prestimulus α Phase Predicts Visual Awareness. J. Neurosci. 29, 2725–2732 (2009).

49. Busch, N. A. & VanRullen, R. Spontaneous EEG oscillations reveal periodic sampling of visual attention. Proc. Natl. Acad. Sci. U. S. A. 107, 16048–16053 (2010).

50. Klimesch, W. EEG-alpha rhythms and memory processes. Int. J. Psychophysiol. Off. J. Int. Organ. Psychophysiol. 26, 319–340 (1997).

51. Bruce, C. J. & Goldberg, M. E. Primate frontal eye fields: I. Single neurons discharging before saccades. J. Neurophysiol. 53(3), 603–635 (1985).

52. Segraves, M. A. & Goldberg, M. E. Functional properties of corticotectal neurons in the monkey’s frontal eye field. J. Neurophysiol. 58, 1387–1419 (1987).

53. Schall, J. D. Neuronal activity related to visually guided saccades in the frontal eye fields of rhesus monkeys: comparison with supplementary eye fields. J. Neurophysiol. 66, 559–579 (1991).

54. Schall, J. D. & Hanes, D. P. Neural basis of saccade target selection in frontal eye field during visual search. Nature 366, 467–469 (1993).

55. Schall, J. D., Hanes, D. P., Thompson, K. G. & King, D. J. Saccade target selection in frontal eye field of macaque. I. Visual and premovement activation. J. Neurosci. Off. J. Soc. Neurosci. 15, 6905–6918 (1995).

56. Schall, J. D. & Thompson, K. G. Neural selection and control of visually guided eye movements. Annu. Rev. Neurosci. 22, 241–259 (1999).

57. Pouget, P. et al. Visual and motor connectivity and the distribution of calcium-binding proteins in macaque frontal eye field: implications for saccade target selection. Front. Neuroanat. 3, 2 (2009).

58. Fries, W. Cortical projections to the superior colliculus in the macaque monkey: a retrograde study using horseradish peroxidase. J. Comp. Neurol. 230, 55–76 (1984).

59. Leichnetz, G. R. & Goldberg, M. E. Higher centers concerned with eye movement and visual attention: cerebral cortex and thalamus. Rev. Oculomot. Res. 2, 365–429 (1988).

60. Sommer, M. A. & Wurtz, R. H. Composition and Topographic Organization of Signals Sent From the Frontal Eye Field to the Superior Colliculus. J. Neurophysiol. 83, 1979–2001 (2000).

61. Buffalo, E. A., Fries, P., Landman, R., Buschman, T. J. & Desimone, R. Laminar differences in gamma and alpha coherence in the ventral stream. Proc. Natl. Acad. Sci. U. S. A. 108, 11262–11267 (2011).

62. Buschman, T. J. & Miller, E. K. Top-down versus bottom-up control of attention in the prefrontal and posterior parietal cortices. Science 315, 1860–1862 (2007).

63. Cavada, C. & Goldman-Rakic, P. S. Posterior parietal cortex in rhesus monkey: II. Evidence for segregated corticocortical networks linking sensory and limbic areas with the frontal lobe. J. Comp. Neurol. 287, 422–445 (1989).

64. Stanton, G. B., Bruce, C. J. & Goldberg, M. E. Topography of projections to posterior cortical areas from the macaque frontal eye fields. J. Comp. Neurol. 353, 291–305 (1995).

65. Corbetta, M. & Shulman, G. L. Control of goal-directed and stimulus-driven attention in the brain. Nat. Rev. Neurosci. 3, 201–215 (2002).

66. Hassen, S. B. H., Gaillard, C., Astrand, E., Wardak, C. & Hamed, S. B. Interneuronal correlations dynamically adjust to task demands at multiple time-scales. bioRxiv 547802 (2019).

67. Tremblay, S., Pieper, F., Sachs, A. & Martinez-Trujillo, J. Attentional filtering of visual information by neuronal ensembles in the primate lateral prefrontal cortex. Neuron 85, 202–215 (2015).

68. Kohn, A., Coen-Cagli, R., Kanitscheider, I. & Pouget, A. Correlations and Neuronal Population Information. Annu. Rev. Neurosci. 39, 237–256 (2016).

69. Ni, A. M., Ruff, D. A., Alberts, J. J., Symmonds, J. & Cohen, M. R. Learning and attention reveal a general relationship between population activity and behavior. Science 359, 463–465 (2018).

70. van Kerkoerle, T. et al. Alpha and gamma oscillations characterize feedback and feedforward processing in monkey visual cortex. Proc. Natl. Acad. Sci. U. S. A. 111, 14332–14341 (2014).

71. Fries, P., Reynolds, J. H., Rorie, A. E. & Desimone, R. Modulation of Oscillatory Neuronal Synchronization by Selective Visual Attention. Science 291, 1560–1563 (2001).

72. Chalk, M. et al. Attention Reduces Stimulus-Driven Gamma Frequency Oscillations and Spike Field Coherence in V1. Neuron 66, 114 (2010).

73. Engel, A. K., Fries, P. & Singer, W. Dynamic predictions: oscillations and synchrony in top-down processing. Nat. Rev. Neurosci. 2, 704–716 (2001).

74. Spyropoulos, G., Bosman, C. A. & Fries, P. A theta rhythm in macaque visual cortex and its attentional modulation. Proc. Natl. Acad. Sci. U. S. A. 115, E5614–E5623 (2018).

75. Ecker, A. S. et al. Decorrelated Neuronal Firing in Cortical Microcircuits. Science 327, 584–587 (2010).

76. Nienborg, H. & Cumming, B. Correlations between the activity of sensory neurons and behavior: how much do they tell us about a neuron’s causality? Curr. Opin. Neurobiol. 20, 376–381 (2010).

77. Trainito, C., von Nicolai, C., Miller, E. K. & Siegel, M. Extracellular Spike Waveform Dissociates Four Functionally Distinct Cell Classes in Primate Cortex. Curr. Biol. 29, R871–R873 (2019).

78. Oostenveld, R., Fries, P., Maris, E. & Schoffelen, J.-M. FieldTrip: Open Source Software for Advanced Analysis of MEG, EEG, and Invasive Electrophysiological Data. Intell Neurosci. 2011, 1:1–1:9 (2011).

79. Grinsted, A., Moore, J. C. & Jevrejeva, S. Application of the cross wavelet transform and wavelet coherence to geophysical time series. Nonlinear Process. Geophys. 11, 561–566 (2004).

80. Buffalo, E. A., Fries, P., Landman, R., Buschman, T. J. & Desimone, R. Laminar differences in gamma and alpha coherence in the ventral stream. Proc. Natl. Acad. Sci. U. S. A. 108, 11262–11267 (2011).

81. Cohen, J. Y., Pouget, P., Heitz, R. P., Woodman, G. F. & Schall, J. D. Biophysical support for functionally distinct cell types in the frontal eye field. J. Neurophysiol. 101, 912–916 (2009).

82. Fiebelkorn, I. C. et al. Cortical cross-frequency coupling predicts perceptual outcomes. NeuroImage 69, 126–137 (2013).

83. Gaillard C., Ben Hadj Hassen S., Di Bello F., Bihan-Poudec Y., VanRullen R., Ben Hamed S. Prefrontal attentional saccades explore space at an alpha rhythm. bioRxiv. doi: https://doi.org/10.1101/637975 (2019)

